# The behavioral effects of gestational and lactational benzo[a]pyrene exposure vary by sex and genotype in mice with differences at the *Ahr* and *Cyp1a2* loci

**DOI:** 10.1101/2021.10.22.465510

**Authors:** Amanda Honaker, Angela Kyntchev, Emma Foster, Katelyn Clough, Emmanuella Asiedu, Mackenzie Feltner, Victoria Ferguson, Philip Tyler Forrest, Jayasree Mullaguru, Mame Diarra Niang, Connor Perry, Yvonne Sene, Christine Perdan Curran

**Author notes:** **Correspondence:** Christine Perdan Curran, Director, Neuroscience Program SC344 Nunn Drive, Department of Biological Sciences, Northern Kentucky University, Highland Heights, KY 41099. These two authors contributed equally to this work (Amanda Honaker, Angela Kyntchev).

## Abstract

Benzo[a]pyrene (BaP) is a polycyclic aromatic hydrocarbon (PAH) and known carcinogen in the Top 10 on the United States’ list of priority pollutants. Humans are exposed through a variety of sources including tobacco smoke, grilled foods and fossil fuel combustion. Recent studies of children exposed to higher levels of PAHs during pregnancy and early life have identified numerous adverse effects on the brain and behavior that persist into school age and adolescence. Our studies were designed to look for genotype and sex differences in susceptibility to gestational and lactational exposure to BaP using a mouse model with allelic differences in the aryl hydrocarbon receptor and the xenobiotic metabolizing enzyme CYP1A2. Pregnant dams were exposed to 10 mg/kg/day of BaP in corn oil-soaked cereal or the corn oil vehicle alone from gestational day 10 until weaning at postnatal day 25. Neurobehavioral testing began at P60 using one male and one female per litter. We found main effects of sex, genotype and treatment as well as significant gene x treatment and sex x treatment interactions. BaP-treated female mice had shorter latencies to fall in the Rotarod test. High-affinity *Ahr^b^Cyp1a2(−/−)* mice had greater impairments in Morris water maze. Interestingly, poor-affinity *AhrdCyp1a2(−/−)* mice also had deficits in spatial learning and memory regardless of treatment. We believe our findings provide future directions in identifying human populations at highest risk of early life BaP exposure, because our model mimics known human variation in our genes of interest. Our studies also highlight the value of testing both males and females in all neurobehavioral studies.

**Highlights:**

- Gestational and lactational benzo[a]pyrene (BaP) exposure has sex and genotype-specific neurobehavioral effects in mice.
- Female mice were more susceptible to motor deficits following developmental BaP exposure. Males were more susceptible to deficits in reversal learning and memory.
- *Ahr^b^Cyp1a2(−/−)* knockout mice were more susceptible to spatial learning and memory deficits following developmental BaP exposure.
- Poor-affinity *AhrdCyp1a2(−/−)* mice had deficits in spatial learning and memory regardless of treatment.

## 1. Introduction

Benzo[a]pyrene (BaP) is a Group 1 carcinogen (IARC 2018) and ranked 8^th^ on the U.S. government’s Priority Pollutants List while the entire class of compounds (polycyclic aromatic hydrocarbons) ranks 9^th^ (ASTDR 2019). Human exposures are widespread from cigarette smoke, wildfires, fossil fuel combustion, and grilled foods (IARC 2018; ASTDR 1996). In addition to cancer risk, there is accumulating evidence from human and animal studies that BaP is neurotoxic and that the developing brain is particularly susceptible to BaP exposure. Chepelev et al. (2015) even suggested that neurotoxicity might be a more sensitive endpoint for human risk assessments than carcinogenicity.

Coke oven workers with high BaP exposure had higher anxiety, impaired reaction time, and more errors on standard tests of neurobehavioral function (Qui et al. 2013). Increased exposure to traffic-related air pollution was associated with a significant decrease in sustained attention in teenagers (Kicinski et al. 2015). Air pollution is also considered a risk factor for autism (Lam et al. 2016) and neurodegenerative disorders including Parkinson’s disease (Lee et al. 2016) and Alzheimer’s disease (Calderón-Garcidueñas et al. 2018).

The most detailed studies including personal backpack monitors, cord blood measurements and long-term assessments of children in New York City found consistent evidence linking developmental PAH exposure to IQ deficits and behavioral problems (Perera et al. 2018; 2014; 2012). Peterson et al. (2015) reported dose-dependent reductions in white matter following both prenatal and postnatal exposure to PAHs. The changes were associated with reduced processing speed, ADHD and other behavioral problems in children tracked up to age 9. There was an inverse relationship between cord blood levels of brain-derived neurotrophic factor (BDNF) and prenatal PAH exposure (Perera et al. 2015). Prenatal PAH exposures have now been linked with academic deficits in adolescents based on tests of language, spelling and math (Margolis et al. 2021). Since many regions with high levels of PAH air pollution also suffer from psycho-social stressors, the adverse outcomes can be magnified for many at-risk populations. For example, Pagliaccio et al. (2020) found a significant interaction between prenatal PAH exposures and early life stress that persisted throughout childhood.

Not all human studies have measured PAH exposure directly, and many rely on other components of air pollution exposure such as particulate matter or nitrogen oxides (Rivas et al. 2019; Min & Min 2017; Sentis et al. 2017; Fuertes et al. 2016). This can make it difficult to pinpoint the specific effects from each type of pollutant, although Jedrychowski et al. (2017) was able to demonstrate that PAH exposure was more important than PM2.5 in reducing birth weights. The number of potential confounding factors highlights the value of animal studies in identifying specific neurotoxic compounds, their mechanism of action and modifying factors such as genetics and nutrition. Indeed, numerous studies have reported neurotoxic effects from BaP exposure in multiple model systems from zebrafish to rats (Lin et al. 2020; Lyu et al. 2020; McCallister et al. 2016)

Our studies were designed to look at genetic differences in the aryl hydrocarbon receptor (AHR) pathway known to alter BaP metabolism (Uno et al. 2006; 2004). We previously found that *Cyp1a2(−/−)* knockout mice were more susceptible to developmental PCB exposure (Colter et al. 2018; Curran et al. 2012; 2011), and maternal CYP1A2 can protect offspring from gestational and lactational exposure to AHR ligands (Dragin et al. 2006; Diliberto et al. 1997). Therefore, we hypothesized that wild-type, high-affinity *Ahr^b^Cyp1a2(+/+)* dams might be better able to sequester and detoxify BaP compared with *Cyp1a2(−/−)* knockouts having either the high-affinity *Ahr^b^* allele or the poor-affinity *Ahr^d^* allele. Using both knockout lines allowed us to further determine if any observed neurotoxicity was AHR-mediated. We used oral dosing to ensure that offspring were exposed only through the placenta and breast milk, because BaP exposure in food can be a significant route of exposure to pregnant women (Wang et al. 2021; Duarte-Salles et al. 2013; Falco et al. 2003) and prior studies indicate oral dosing is the preferred route of exposure for assessing gene-environment interactions related to BaP toxicity (Nebert et al. 2013).

## 2. Methods

### 2.1 Animals

All experiments and care were conducted under protocols approved by the Northern Kentucky University Institutional Animal Care and Use Committee (IACUC) and in accordance with the ARRIVE Guidelines (Percie du Sert et al. 2020; Kilkenny et al., 2010). Animal care and housing was in accordance with the Guide for the Care and Use of Laboratory Animals (8^th^ ed.).

#### 2.1.1. Animal source and husbandry

Male and female C57BL/6J mice were purchased from The Jackson Laboratory (Bar Harbor, ME). Animals were group housed with a maximum of 4 per cage by sex and treatment group in standard polysulfone shoebox cages with static microfilter lids, corncob bedding and a 12 h:12 h light-dark cycle. Water and Lab Diet 5015 chow (18.9% protein; 11% fat, 32.6% starch) were provided *ad libitum*. Animals were checked daily for health concerns, and cages were changed weekly with one cotton nestlet provided for enrichment. Animals were acclimated to the vivarium for >1 wk prior to beginning experimental treatments. *Cyp1a2(−/−)* knockout mice were originally back-crossed at least eight generations onto a B6 background. Colonies have been maintained in the Northern Kentucky University vivarium, and knockout lines were routinely back-crossed to the B6 breeders from The Jackson Laboratory to avoid confounding by genetic drift. Nulliparous females were mated with males of the same genotype on a four-day breeding cycle. The presence of a vaginal plug was considered evidence of mating, and dams were removed from the breeding cage at that time. Litters were culled or cross-fostered to balance litter size at six pups per dam.

#### 2.1.2. Benzo[a]pyrene treatments

Benzo[a]pyrene (> 96% purity) was purchased from Millipore-Sigma and dissolved in corn oil. At gestational day 10, pregnant dams were randomly assigned to receive either 10/mg/kg/day BaP in corn oil-soaked cereal (Cap’n Crunch peanut butter) or the corn oil vehicle until pups were weaned at postnatal day 25 (P25). The dose was based on our preliminary studies to avoid overt toxicity and similar rodent studies designed to mimic known human exposures (Chen et al. 2012; Zhang et al. 2016) while accounting for the higher metabolic rate of laboratory mice.

#### 2.1.3. Verification of genotype

Tail snips (~ 1 cm) were taken, and PCR was used to verify the genotypes of all dams, sires and offspring used in behavioral tests. Data from animals not matching the expected genotype was excluded from analysis (n=4).

### 2.2 Behavioral testing

To verify that offspring received equivalent care, we used two tests of maternal behavior during the first week following birth. One male and one female pup from each litter were randomly selected at P25 for behavioral testing that began at P60. Experiments are presented in the order in which they were conducted. Mice underwent no more than one behavioral test per day during the light cycle, and testing was restricted to a four-hour time block to avoid confounding by circadian rhythms. Personnel handling animals for behavior experiments were blinded to treatment groups. Lighting was set at 50% of full brightness in all rooms used for behavioral testing. Male and female mice were tested separately on each apparatus.

#### 2.2.1 Dam behavior

We conducted snapshot ethograms of dam behavior within 1 h of lights on and 1 h of lights off to assess general care of pups and maternal behavior. The percent time spent on nest during each 1 h observation period was compared. We also used a scale of 0-3 to rate nest quality with 0 being the absence of a nest and 3 being a fully developed nest including roof.

#### 2.2.2 Open field locomotor activity

Mice were placed in a square plexiglass chamber (41 cm x 41 cm) for 60 min while a Photobeam Activity System (San Diego Instruments, San Diego, CA) tracked animal activity during 12 five-min intervals. The number of beam breaks was used as a measure of ambulatory activity. A second row of photobeams detected rearing movements. Chambers were cleaned with 70% ethanol between each test.

#### 2.2.3 Acoustic startle with pre-pulse inhibition

The SR-Lab apparatus (San Diego Instruments, San Diego, CA) was used to test baseline startle response and sensorimotor gating with a 120 db startle stimulus and pre-pulses of 74 and 76 db. A Latin Squares design was used with four trial types randomly presented over a total of 48 trials. Peak response amplitudes (Vmax) were recorded, and the percent attenuation following the pre-pulse tones was calculated (Curran et al. 2012).

#### 2.2.4 Pole climb

To test motor coordination, mice were placed at the top of a 50 cm pole facing upwards. The time required to turn downward and total time to descend to the home cage were recorded (Colter et al. 2018).

#### 2.2.5 Rotarod

After two days of acclimation, mice were tested for five days on a Rotarod (IITC Life Science Inc., Woodland Hills, CA.) to assess motor coordination and motor learning. The Rotarod was programmed to accelerate from 0-20 rpm over 180 s with a 300 s maximum trial time. Mice were given three trials/day and the latency to fall or slip was recorded. The apparatus was cleaned between each trial to avoid odor interference (Colter et al. 2018).

#### 2.2.6 Novel object recognition

Mice were acclimated to a circular arena (91 cm) with no objects present for two days followed by two days of acclimation with objects present similar in size to test objects, but with distinctly different shapes and colors. On the fifth day, mice accumulated 30 s of exploration of two identical objects in the familiarization phase. One hour later, visual recognition memory was tested by presenting a copy of the familiar object and a novel test object. Again, each mouse accumulated 30 s total exploration time. The percent time exploring the novel object was compared. Familiar and test objects were counter-balanced to avoid confounding by object preference (Brown et al. 2020).

#### 2.2.7 Morris water maze

The Morris water maze was used to assess hippocampal dependent spatial learning and memory using a four-phase paradigm. (Brown et al. 2020, Curran et al., 2011, 2012; Vorhees and Williams, 2006). In each phase, mice were placed in random start locations around a 112 cm tank with the goal of finding an escape platform. Water temperature was maintained at 22° C ± 2° C, and the water was colored with non-toxic white tempera paint to obscure the platform location. In the Cued phase, curtains were drawn and an orange ping-pong ball on a pole in the center of a 10 cm circular platform served as a proximal cue to the escape platform’s location. The Cued phase demonstrated that mice could swim and could use visual cues to find the escape platform. All mice successfully completed this phase and moved into the Hidden Platform phases. Mice received 4 trials/day with a 15 min inter-trial interval for 6 days followed by a 30 s Probe trial with the platform removed on day 7. Curtains were opened to reveal distal cues around the room. Platform size decreased each week from 10 cm to 7 cm to 5 cm, and each week the platform was moved to a new location. Latency to escape, distance traveled, swim speed, and average distance to the escape platform were recorded by AnyMaze™ software (Stoelting, Inc.). Mice that did not reach the platform within 60 s were guided to the platform and kept on the platform for 15 s. For probe trials, average distance to the target and target zone crossings were recorded.

#### 2.2.8 Data analysis

Data were analyzed by the KY-INBRE Applied Statistics Laboratory at the University of Kentucky using SAS Proc Mixed and slice effects when significant differences were found. Corrections were made for multiple post-hoc analyses. Swim speed was used as a covariate with latency to escape in Morris water maze. A repeated measures design was used for Rotarod and Morris water maze with day as the repeated measure and in Open Field Locomotor with interval as the repeated measure. Significance was set as P < 0.05, and trends are noted if P < 0.1. Data are presented as LS means, and error bars represent the standard error of the mean. Based on prior experience with mouse behavior, we used 15-20 litters per group to generate sufficient power for analyses.

## 3. Results

### 3.1 Dam behavior

There were no significant differences in maternal care based on genotype or treatment when comparing time spent on nest and nest quality (P > 0.05; data not shown).

### 3.2 Open field locomotor activity

There was a significant gene x treatment interaction when comparing time spent in the periphery v. the central region of the arena. BaP-treated *Ahr^d^Cyp1a2(−/−)* mice spent less time in the central region, which is generally considered an indication of increased anxiety-like behavior (P < 0.05; Fig. 1). There was also a main effect of genotype with *Ahr^b^Cyp1a2(−/−)* mice spending significantly more time in the central region compared with the other two genotypes. When comparing total beam breaks in each 5-minute interval, BaP-treated high-affinity *Ahr^b^Cyp1a2(−/−)* mice were more active than their corn oil controls while BaP-treated poor-affinity *Ahr^d^Cyp1a2(−/−)* mice were less active than their corn oil controls. However, the differences were only significant in one of the 12 intervals. All groups showed normal habituation over the 1 h period with lower activity in the later intervals (data not shown).

**Fig. 1.**
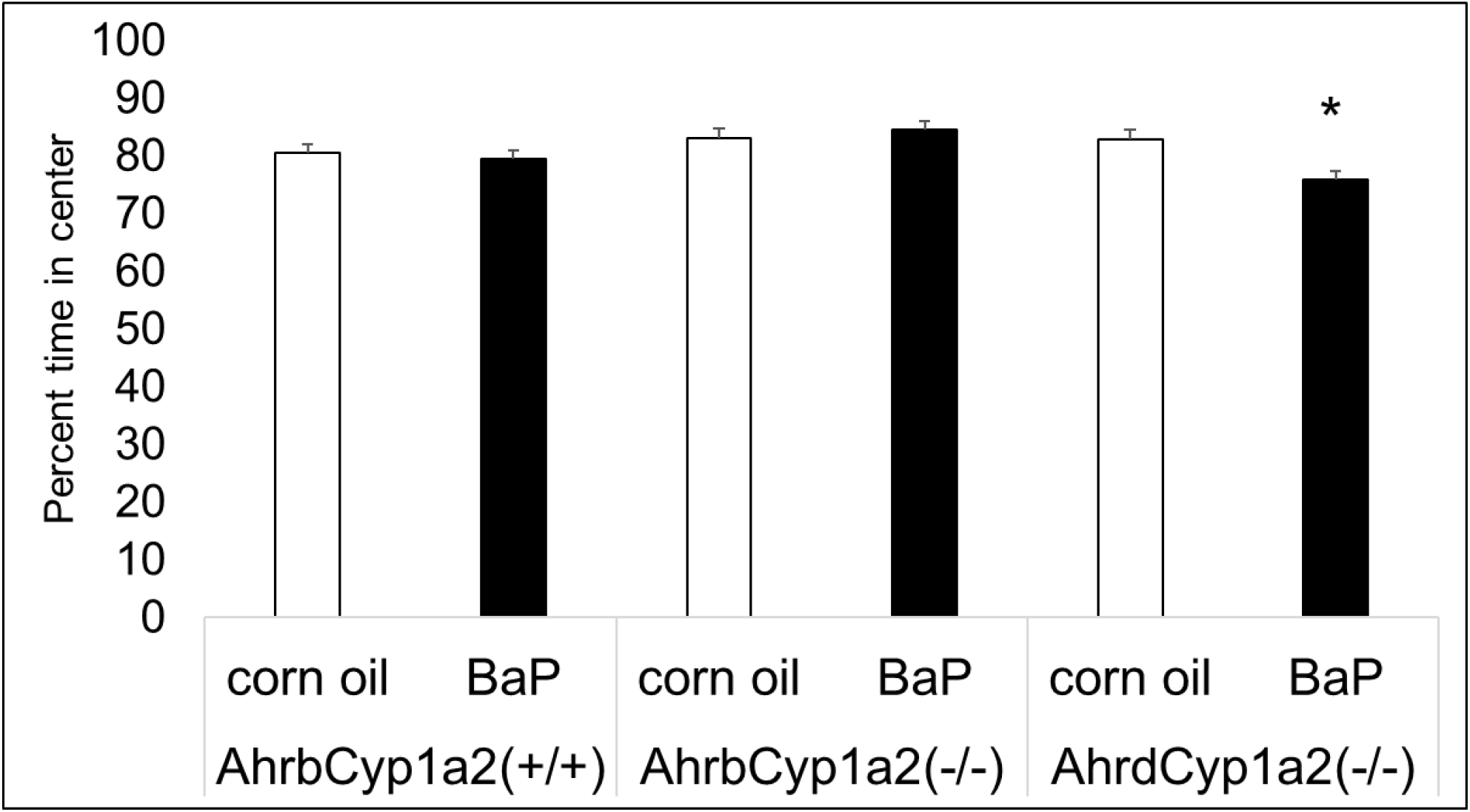
Open field locomotor activity. BaP-exposed *Ahr^d^Cyp1a2(−/−)* mice spent significantly less time in the central area of the apparatus compared with all other groups. * P < 0.05

### 3.3 Acoustic startle with pre-pulse inhibition

We found the expected sex difference in the baseline startle response with heavier males having a higher amplitude compared with females (P < 0.001) and a significant main effect of genotype with high-affinity *Ahr^b^Cyp1a2(−/−)* mice having a much lower baseline startle response compared with the other two genotypes (P < 0.001; Fig.2). There was no effect of treatment or genotype on sensorimotor gating with all groups showing attenuation under the two pre-pulse conditions.

**Fig. 2.**
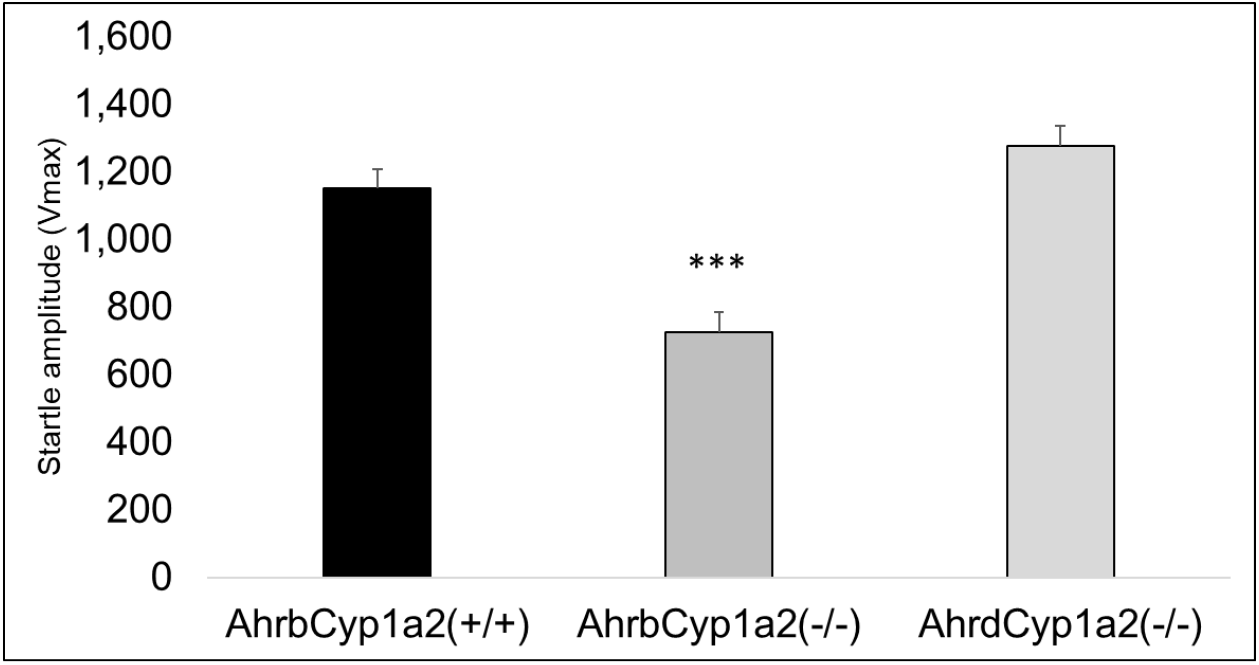
Baseline startle response. *Ahr^b^Cyp1a2(−/−)* mice had a significantly lower response to the 120 db startle stimulus compared with the other two genotypes. *** P < 0.001

### 3.4 Pole climb

There was a main effect of genotype in the pole climb experiment. High-affinity *Ahr^b^Cyp1a2(−/−)* mice had significantly shorter latencies to turn and to descend the pole (P < 0.001; Fig. 3). There was also a main effect of sex with females having shorter latencies to turn (P < 0.05) and descend (P < 0.01), but there was no effect of BaP treatment (P > 0.05; data not shown).

**Fig. 3.**
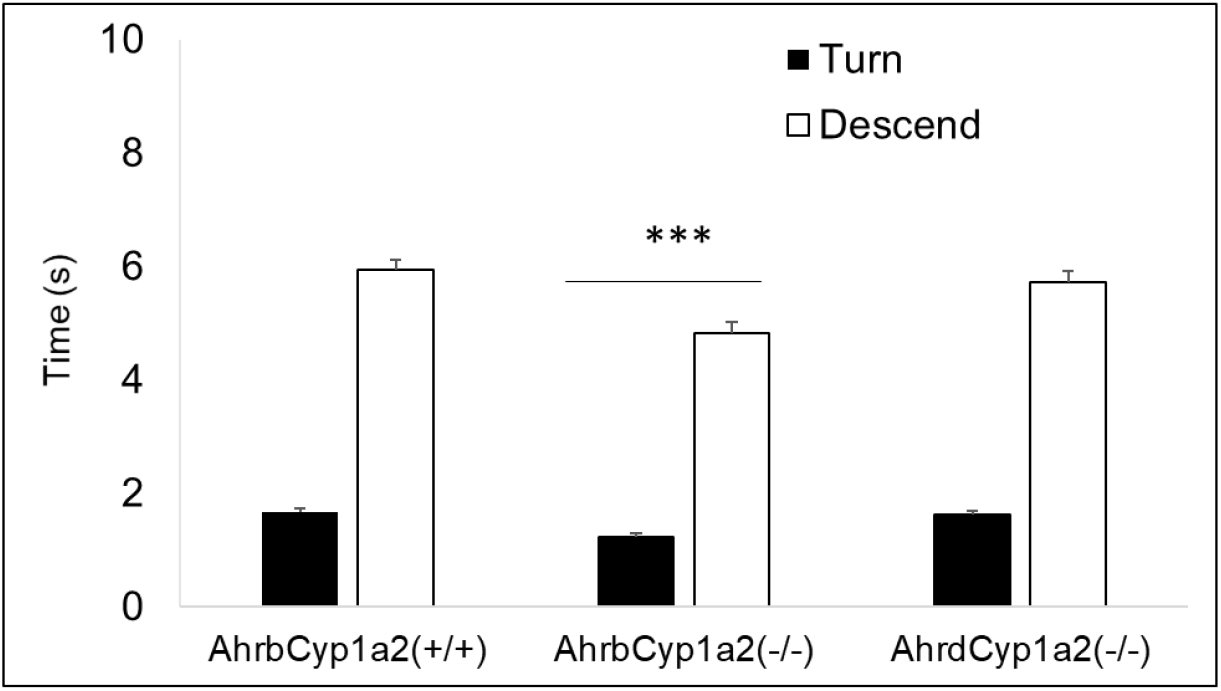
Pole climb results. *Ahr^b^Cyp1a2(−/−)* mice had a significantly faster times to turn and descend a 50 cm pole. *** P < 0.001

### 3.5 Rotarod

There was a significant sex x treatment interaction in the Rotarod test. BaP-treated female mice had shorter latencies to fall off the rotarod on all five days of testing compared with control females; however, the differences were only significant on two of five days of testing (P < 0.05; Fig. 4A). Interestingly, BaP-treated males had longer latencies than corn oil-treated males on days 3-5 of testing and significantly longer latencies on two days of testing indicating greater motor learning over the 5-day test (Fig. 4B).

**Fig. 4A-B.**
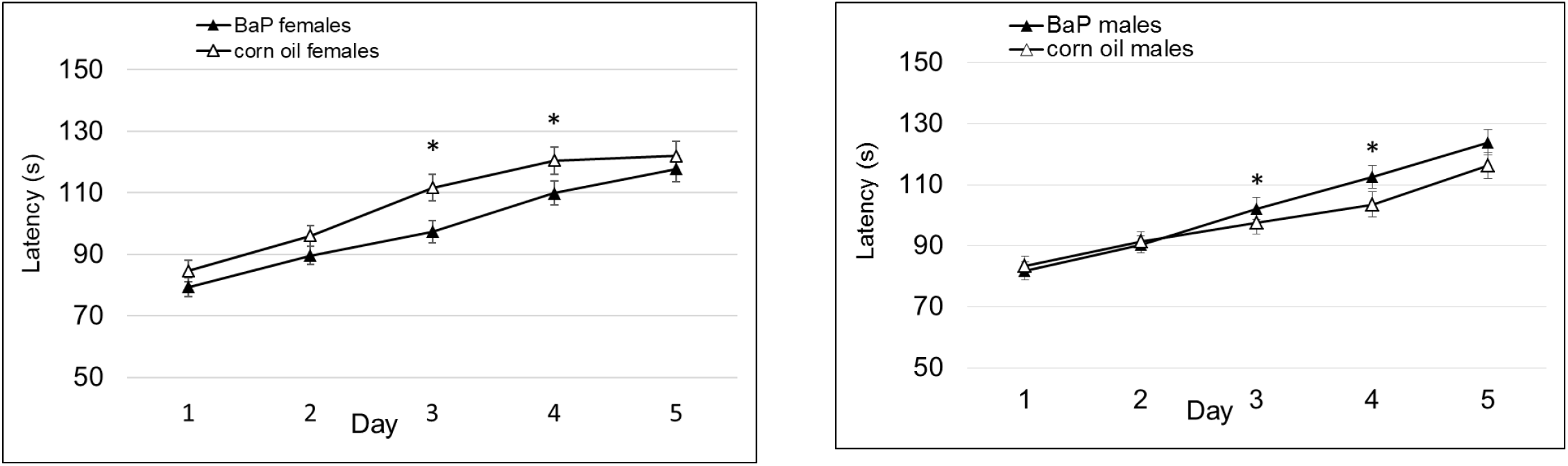
Rotarod results. **A.**BaP-exposed female mice had shorter latencies to fall off the rotarod on days 3 and 4 of testing. **B.** BaP-exposed males had longer latencies on days 3 and 4. * P < 0.05

### 3.6 Novel object recognition

There was a trend for significance in Novel Object Recognition with BaP-treated mice spending less time exploring the novel object (P = 0.095; Fig. 5A). Interestingly, the greatest differences were seen in the two lines of *Cyp1a2(−/−)* knockout mice although there was not a significant gene x treatment interaction (P > 0.05; Fig. 5B).

**Fig. 5A-B.**
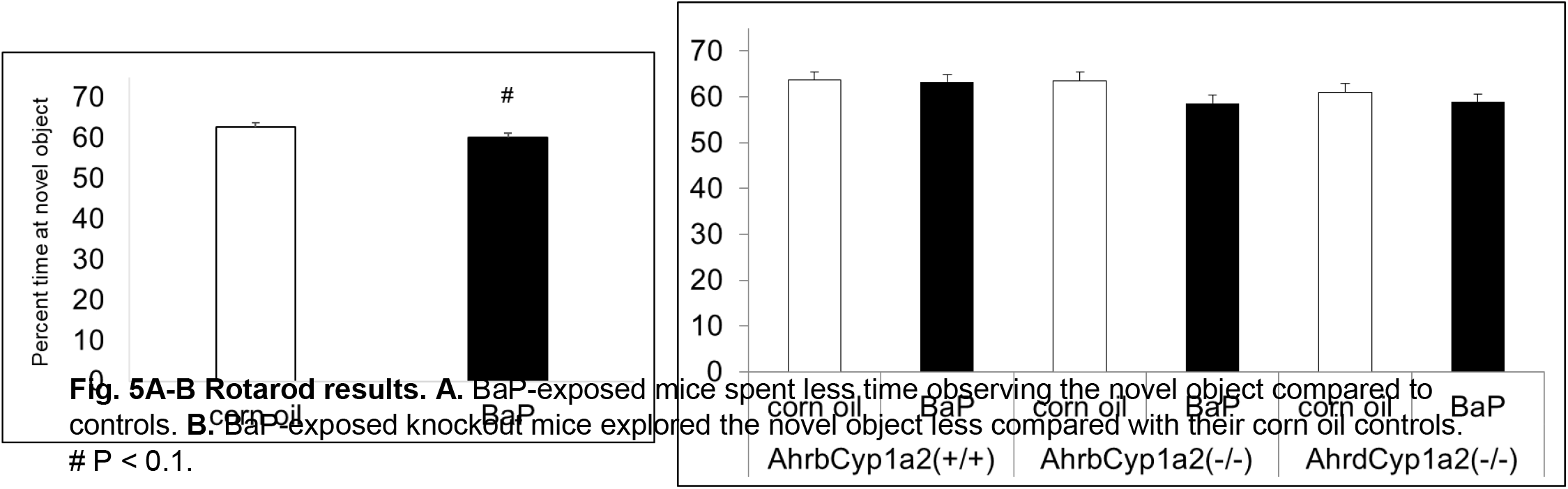
Rotarod results. **A.** BaP-exposed mice spent less time observing the novel object compared to controls. **B.** BaP-exposed knockout mice explored the novel object less compared with their corn oil controls. # P < 0.1.

### 3.7 Morris water maze

In the Cued phase with a visible cue on the platform, all mice were able to swim and locate the escape platform. However, poor-affinity *Ahr^d^Cyp1a2(−/−)* mice had significantly longer escape latencies on four of six days of testing (Fig. 6). There was a main effect of treatment (P < 0.05) on day2 and 6 with BaP-treated mice having longer latencies compared with corn oil-treated controls.

**Fig.6.**
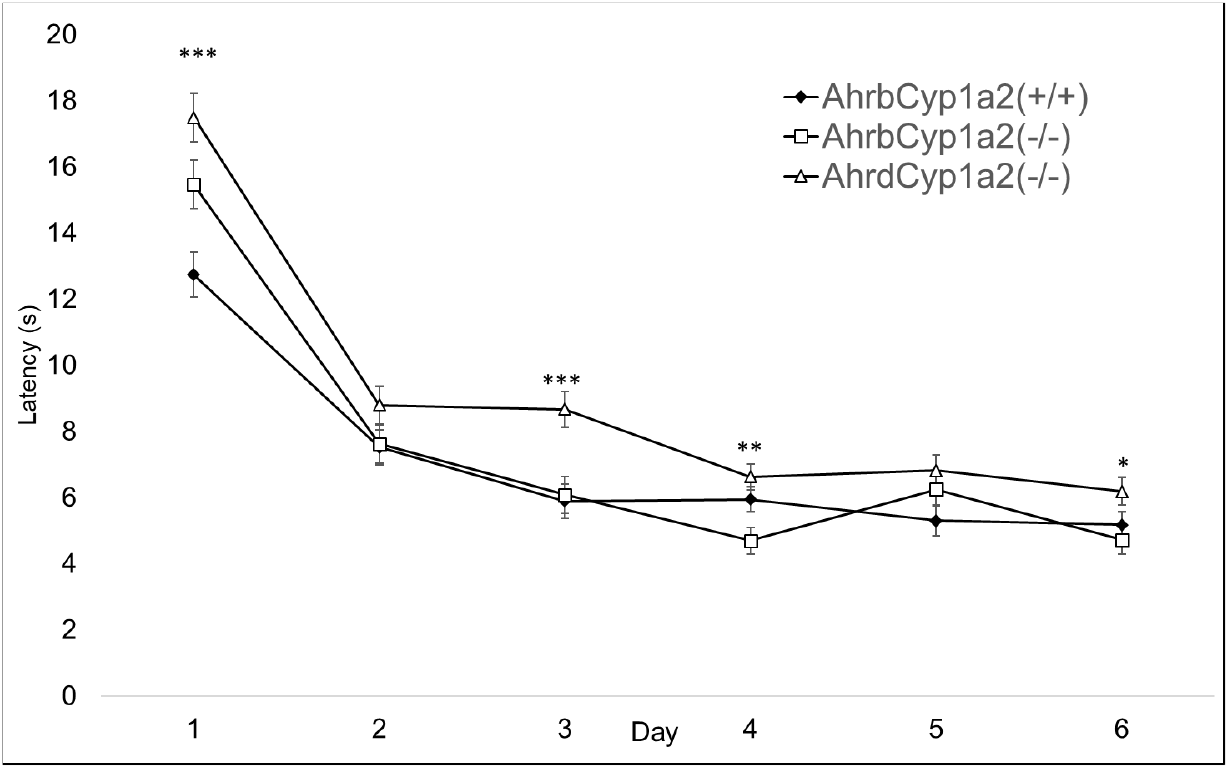
Morris water maze Cued Phase. Poor affinity *Ahr^d^Cyp1a2(−/−)* knockout mice had longer latencies to escape on days 1, 3, 4 and 6. * P < 0.05, ** P < 0.01, *** P < 0.001

In the hidden platform phases, there was a significant main effect of genotype in all three phases with poor-affinity *Ahr^d^Cyp1a2(−/−)* mice having longer path lengths to the escape platform on all but two days of testing (Figs 7A-C). In the Acquisition phase of the hidden platform trials, there was a main effect of treatment on three of the first four days of testing, indicating it took BaP-treated mice longer to learn the location of the escape platform. There was a significant gene x treatment interaction on two days of testing and a trend toward significance on two other days with BaP-treated *Ahr^b^Cyp1a2(−/−)* knockouts showing the greatest impairments (Figs. 8A-C). In the Reverse phase, there was a significant main effect of treatment on four days of testing with BaP-treated mice having longer average distances to the escape platform (Fig. 9) and a significant sex x treatment interaction (P < 0.05) on two days of testing with BaP-treated males having longer path lengths compared to corn oil-treated control males. The opposite trend was seen in females. There was a trend for significance of treatment on Day 5 of the Shift-reduced phase (P = 0.051) and a significant difference on Day 6 for average distance to the platform (P < 0.01).

**Fig. 7A-C.**
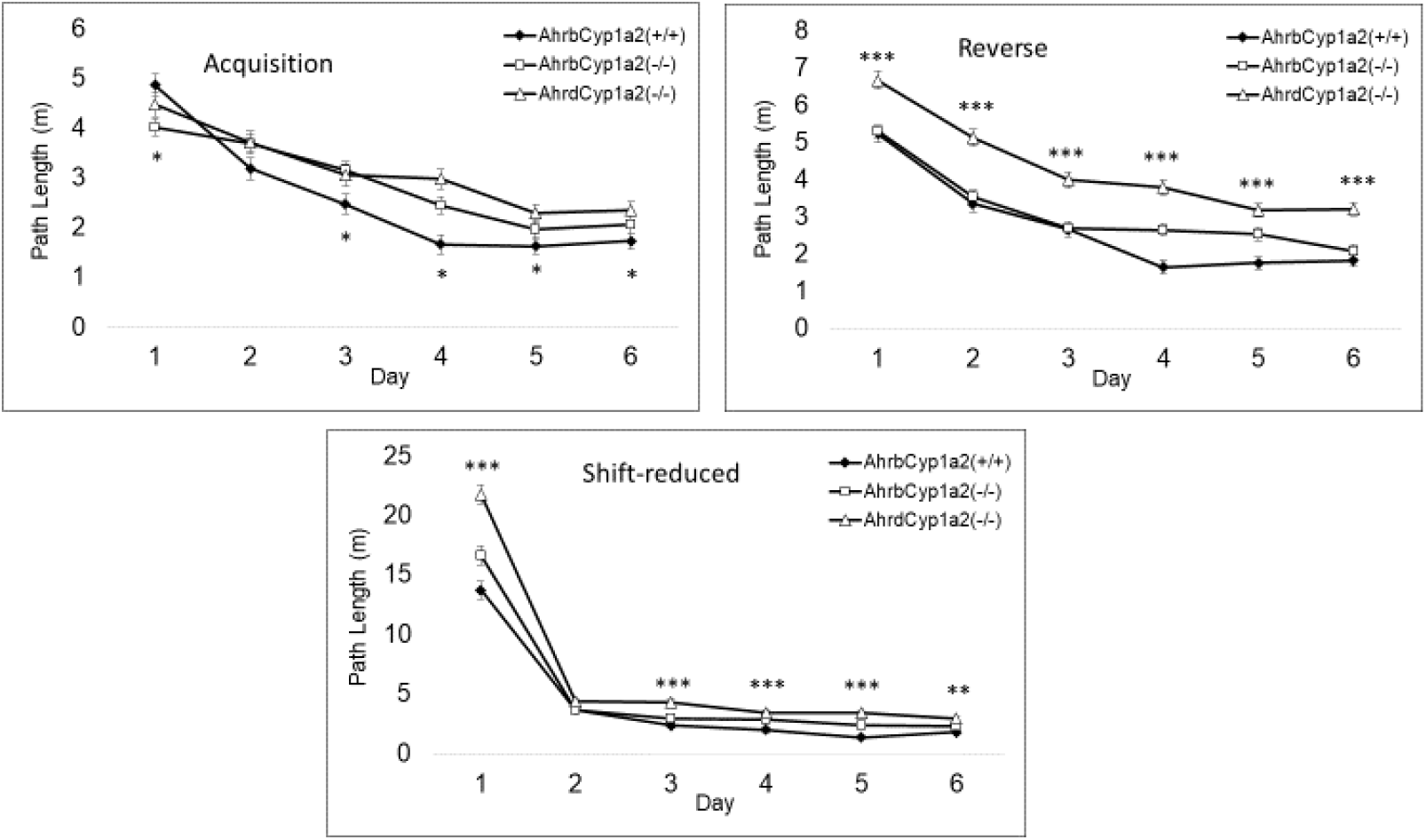
Morris water maze Hidden Phases. Poor affinity *Ahr^d^Cyp1a2(−/−)* knockout mice had longer path lengths on days 1, 3, 4, 5 and 6 in the Acquisition phase (A) on all six days of testing in the Reverse phase (B) and on days 1, 3, 4, 5 and 6 of the Shift-reduce phase (C). * P < 0.05, ** P < 0.01, *** P < 0.001

**Fig. 8A-C.**
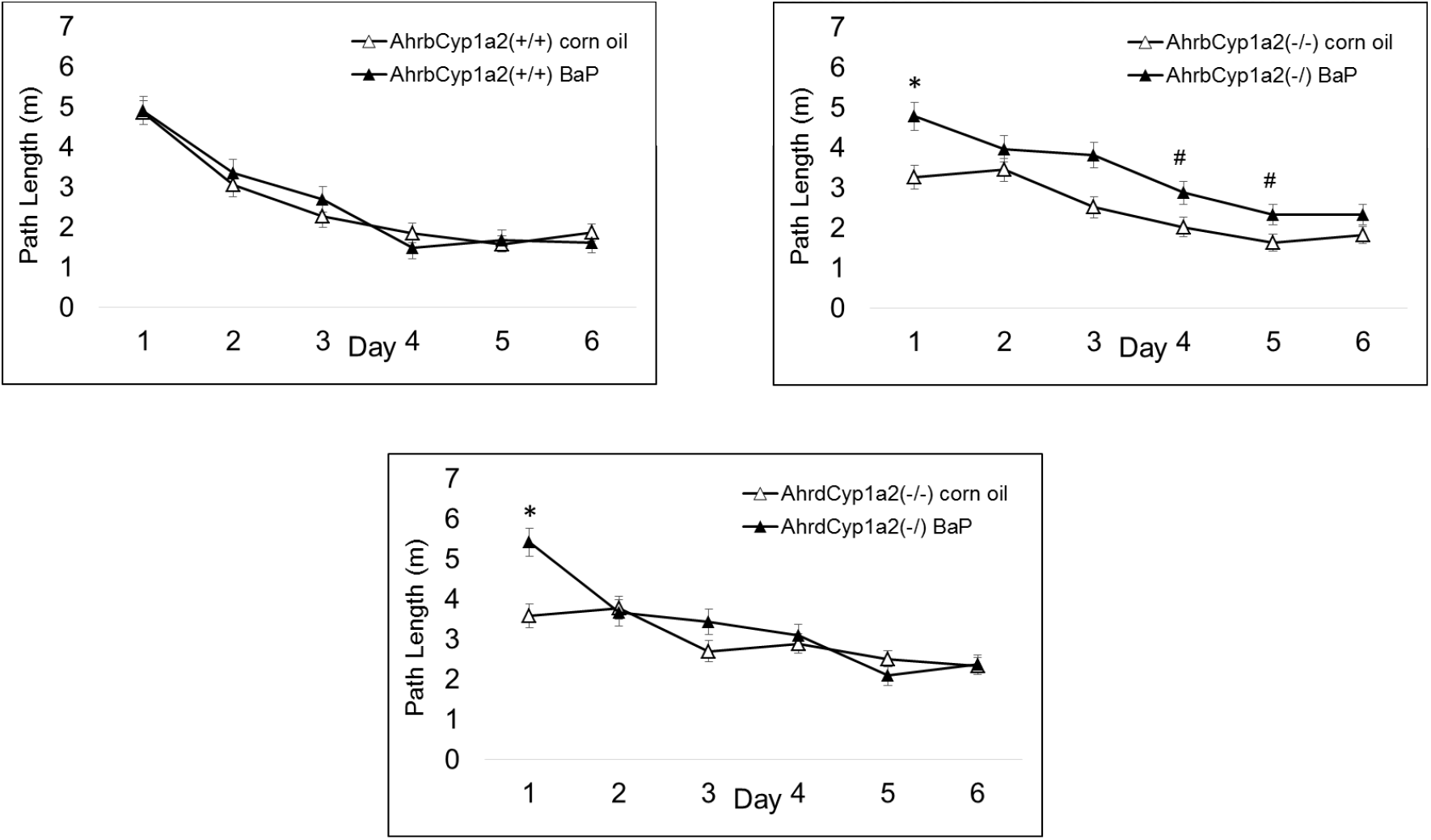
Morris Acquisition gene x treatment interaction. BaP-exposed high-affinity *Ahr^b^Cyp1a2(−/−)* knockout mice had significantly longer path lengths on days 1 and a trend for significance on days 4 and 5. # P < 0.1, * P < 0.05

**Fig. 9.**
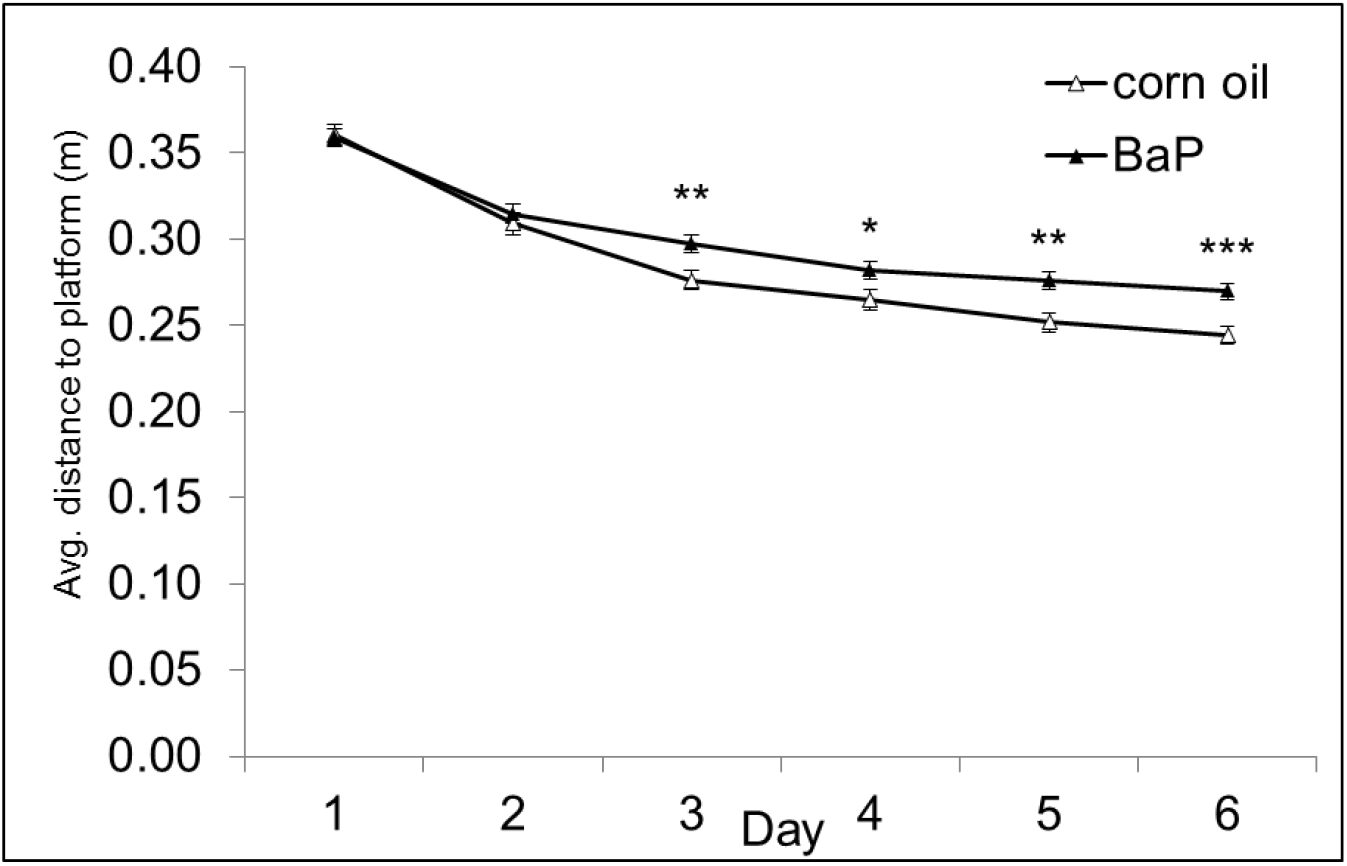
Morris Reverse treatment effect. BaP exposed mice had significantly longer average distances to the escape platform on days 3-6. * P < 0.05, ** P < 0.01, *** P < 0.001

In the Probe trials, BaP- treated mice had significantly longer average distances to the platform in the Acquisition and Reverse phases (P < 0.05) and a trend for significance in the Shift-reduced phase (P = 0.68, Fig.10). There was a significant gene x treatment interaction for target zone crossings in the Shift-reduced phase with BaP-treated *Ahr^b^Cyp1a2(+/+)* wild type mice having more crossings than their corn oil-treated controls. In contrast, BaP-treated *Ahr^b^Cyp1a2(−/−)* knockout mice had fewer crossings than their corn oil-treated controls (P < 0.05).

**Fig. 10.**
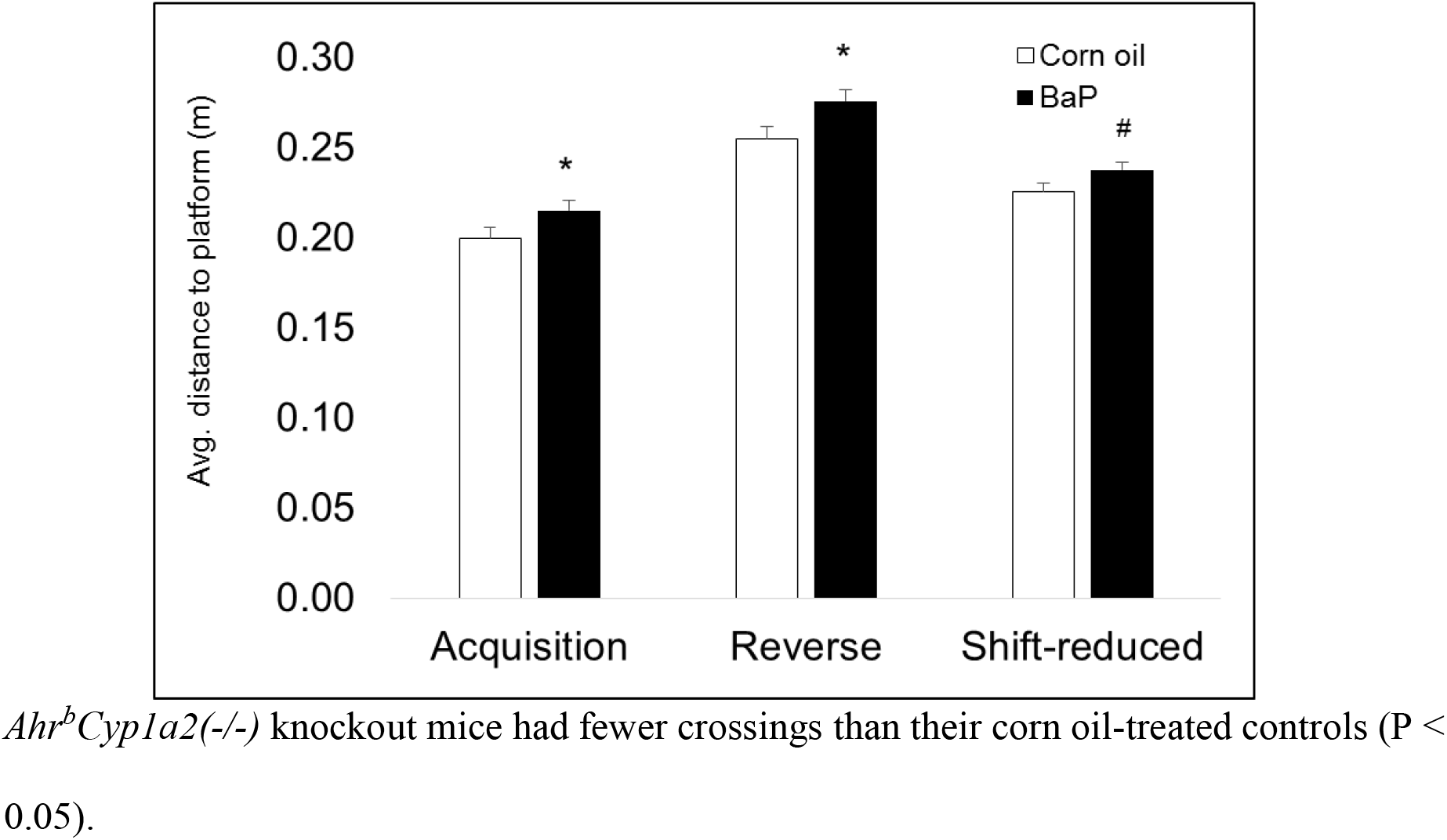
Morris Probe trials. BaP exposed mice had significantly longer average distances to the platform location in the Acquisition and Reverse probe trials and a trend for significance in the Shift-reduced phas. * P < 0.05, # P < 0.1

## 4. Discussion

Using a mouse model with allelic differences at the *Ahr* and *Cyp1a2* loci, we examined adult offspring exposed to benzo[a]pyrene (BaP) during gestation and lactation, a time of critical brain development. We found a variety of treatment, sex and genotype effects and multiple interactions using a battery of behavioral tests to assess locomotor activity, sensorimotor gating, motor function, and learning and memory. Our findings of BaP neurotoxicity are consistent with previous rodent studies (Patel et al. 2016; McCallister et al. 2016; Maciel et al. 2014). Importantly, there were no differences in maternal care based on two measures of dam behavior. Therefore, it is highly likely that the adverse effects reported here were due to BaP exposure and the reported interactions of genotype x treatment and sex x treatment have helped to identify the groups most susceptible to early life BaP exposure.

BaP-treated poor-affinity *Ahr^d^Cyp1a2(−/−)* knockout mice spent significantly less time in the central area of the open field locomotor apparatus compared with all other groups; however, the percentage of time in the center was still well above 70%. Therefore, it would be difficult to conclude that this indicates increased anxiety-like behavior. There was no effect of BaP treatment in the Pole Climb test, but there was a sex x treatment interaction in Rotarod. BaP-treated females were impaired relative to corn oil-treated females whereas BaP-treated males out-performed corn oil-treated males overall. Liu et al. (2002) and Patel et al. (2016) both reported impairments in Rotarod following BaP treatment in male Wistar rats, but to our knowledge, this is the first report of sex-specific effects in Rotarod following developmental BaP exposure. In human studies, Dix-Cooper et al. (2012) reported impaired fine motor skills in school-aged children exposed to wood smoke in cooking fires and carbon monoxide. Our findings indicate that PAH exposure should also be considered as a possible causative agent.

It will also be important to look for sex differences in BaP tissue levels. Singh et al. (1997) demonstrated that higher expression of three glutathione S-transferases in male A/J mice protected them from tumors following BaP exposure compared with females. Although all mice in this study are on a C57BL/6J background, it’s plausible that these females were also less able to detoxify BaP, resulting in a higher body burden and greater neurotoxicity compared with males. On the other hand, there was a sex x treatment interaction in the Reverse phase of Morris water maze with BaP-exposed males having greater impairments than females. Therefore, future studies should include neurotransmitter analyses of all relevant brain regions to determine if there are sex-dependent effects that could explain the observed differences in susceptibility.

BaP-exposed high-affinity *Ahr^b^Cyp1a2(−/−)* knockout mice had longer path lengths compared to corn oil control mice in the Reverse Phase of Morris water maze, and both lines of BaP-exposed knockout mice spent less time exploring the novel object compared with wild type mice. This indicates that both *Cyp1a2* and *Ahr* genotype are important when considering susceptibility to developmental BaP exposure on measures of hippocampal dependent learning and memory. Both genes are highly variable in the human population (Zhou et al. 2010; Nebert 2017; Nebert et al. 2013), and our mouse model was designed to mimic that genetic variation. Further work is needed to determine how genotype affects the transfer of BaP from dam to fetus and offspring during gestation and lactation, but previous work with AHR agonists clearly indicate that maternal genotype is critical to sequestering and metabolizing persistent organic pollutants such as coplanar PCBs and dioxins (Curran et al. 2011; Dragin et al. 2006; Diliberto et al. 1997). CYP1A1 induction will be higher in *Ahr^b^* mice compared with *Ahr^d^* mice, and both CYP1A1 and CYP1A2 are expressed in liver and the intestines (Uno et al. 2008). Therefore, it is likely we will find different levels of the parent BaP compound and metabolites in tissues from each genotype tested and that differential toxicokinetics could account for the observed differences in neurotoxicity. This is important, because Patel et al. (2016) reported that acute neonatal exposure to BaP increased oxidative stress in the hippocampus, a region essential for normal learning and memory. We would expect to see higher levels of oxidative stress in animals with a higher body burden during early brain development, which would suggest a potential mechanism for differential neurotoxicity.

Beyond the BaP effects, we found poor-affinity *Ahr^d^Cyp1a2(−/−)* knockout mice had significant impairments in the Cued and Reverse phases of Morris water maze. This is consistent with our previous findings of impaired spatial learning and memory in this line of mice (Curran et al. 2012). High-affinity *Ahr^b^Cyp1a2(−/−)* knockout mice had shorter latencies to turn and descend in the Pole Climb test. Although this cannot be considered an impairment, it does suggest a difference in motivation or potentially anxiety-like behavior that can be further explored with tests such as Zero Maze and Marble Burying. These results emphasize the importance of thoroughly characterizing the neurobehavioral phenotype of knockout mice prior to drawing conclusions relative to environmental exposures. Given recent findings regarding the role of AHR activation and the microbiome (Korecka et al. 2016; Nebert 2017) and the effects of BaP on the gut microbiome (Ribière et al. 2016), these mice are likely to be extremely useful in understanding normal and perturbed signaling along the gut-brain axis.

Given the widespread of exposure to BaP through fossil fuel combustion, wildfires, and grilled food (ASTDR 2019; Jedrychowski et al. 2012), the impact of PAH exposure on human health will remain an important public health concern. Recent studies have strongly associated prenatal and early life PAH exposure with adverse behavioral and neurological outcomes in school-aged children and adolescents (Margolis et al. 2021; Perera et al. 2018; Peterson et al. 2015). Teasing out the causative agent and threshold for adverse effects becomes more difficult when studying human populations because there are typically multiple co-exposures such as ozone, nitrogen oxides and particulate matter. Individuals living in urban areas with high levels of traffic-related air pollution are also more likely to face additional psycho-social stressors and the lingering contamination from lead paint and lead water lines.

## 5. Conclusions

We found expected BaP-mediated impairments in learning and memory with high-affinity *Ahr^b^Cyp1a2(−/−)* showing the greatest susceptibility to developmental BaP exposure. We identified sex as a key variable for motor function deficits with BaP-exposed females having impairments whereas males did not. Intriguingly, we found numerous differences based on genotype alone. These findings replicate and extend our previous work with *Cyp1a2(−/−)* knockout mice and suggest that our genes of interest have a normal function in brain development or function. This emphasizes the importance of using intact animal models to gain a deeper understanding of neurodevelopment and neurobehavior and the potential value of these lines of research.

## 6. Author contributions

All authors made substantial contributions to the experimental work and to the development of the manuscript. The two first authors prepared the draft manuscript and assisted with treatments, animal care, data collection and preparation of data for statistical analysis. The other co-authors all assisted with treatments, animal care, neurobehavioral testing and review of the manuscript. The senior author Curran served as the principal investigator responsible for training, experimental design and oversight of the project.

## 7. Acknowledgements

We acknowledge support from multiple funding sources including NIH grants R15ES030541, R15ES020053, P20GM103436 and NSF RSF-034-07, Society of Toxicology internship grants, and the following Northern Kentucky University sources: College of Arts and Sciences Collaborative Faculty-Student Project Awards, Faculty Development Project Grants, Center for Integrative Natural Sciences and Mathematics (CINSAM) Research Grants, CINSAM UR-STEM Fellowships, Greaves and Herrmann Fellowships, and Student Undergraduate Research and Creative Activities awards. We thank Dr. Daniel W. Nebert (University of Cincinnati Medical Center) for the generous donation of *Cyp1a(−/−)* knockout mice. We gratefully acknowledge statistical assistance from Melinda MacDougall. Finally, the encouragement, guidance and wisdom of Dr. Philip J. Bushnell must be acknowledged as well as his legacy of fostering the highest level of rigor and reproducibility in behavioral toxicology.

## 8. Conflicts of interest

The authors have no conflicts to declare.

## 9. Data statement

All data are available upon request as well as detailed protocols for all experiments described herein. Portions of these data were presented previously at the annual meetings of the Society of Toxicology, the Developmental Neurotoxicology Society and the Society for Birth Defects Research and Prevention.

## Notes

### Competing Interest Statement

The authors have declared no competing interest.

## Literature Cited

ATSDR (Agency for Toxic Substances and Disease Registry). 1995. Toxicological profile for polycyclic aromatic hydrocarbons. Atlanta, GA: U.S. Department of Health and Human Services, Public Health Service.

ATSDR (Agency for Toxic Substances and Disease Registry). 2018. Substance Priority List. https://www.atsdr.cdc.gov/spl/index.html#2019spl

Brown, J., Villalona, Y., Weimer, J., Ludwig, C.P., Hays, B.T., Massie, L., Marczinski, C.A., Curran, C.P., 2020. Supplemental taurine during adolescence and early adulthood has sex-specific effects on cognition, behavior and neurotransmitter levels in C57BL/6J mice dependent on exposure window. Neurotoxicol Teratol 79, 106883. https://doi.org/10.1016/j.ntt.2020.106883

Calderón-Garcidueñas, L., Gónzalez-Maciel, A., Reynoso-Robles, R., Delgado-Chávez, R., Mukherjee, P.S., Kulesza, R.J., Torres-Jardón, R., Ávila-Ramírez, J., Villarreal-Ríos, R., 2018. Hallmarks of Alzheimer disease are evolving relentlessly in Metropolitan Mexico City infants, children and young adults. APOE4 carriers have higher suicide risk and higher odds of reaching NFT stage V at ≤40 years of age. Environ Res 164, 475–487. https://doi.org/10.1016/j.envres.2018.03.023

Chen, C., Tang, Y., Jiang, X., Qi, Y., Cheng, S., Qiu, C., Peng, B., Tu, B., 2012. Early postnatal benzo(a)pyrene exposure in Sprague-Dawley rats causes persistent neurobehavioral impairments that emerge postnatally and continue into adolescence and adulthood. Toxicol Sci 125, 248–261. https://doi.org/10.1093/toxsci/kfr265

Chepelev, N.L., Moffat, I.D., Bowers, W.J., Yauk, C.L., 2015. Neurotoxicity may be an overlooked consequence of benzo[a]pyrene exposure that is relevant to human health risk assessment. Mutat Res Rev Mutat Res 764, 64–89. https://doi.org/10.1016/j.mrrev.2015.03.001

Colter, B.T., Garber, H.F., Fleming, S.M., Fowler, J.P., Harding, G.D., Hooven, M.K., Howes, A.A., Infante, S.K., Lang, A.L., MacDougall, M.C., Stegman, M., Taylor, K.R., Curran, C.P., 2018. Ahr and Cyp1a2 genotypes both affect susceptibility to motor deficits following gestational and lactational exposure to polychlorinated biphenyls. Neurotoxicology 65, 125–134. https://doi.org/10.1016/j.neuro.2018.01.008

Curran, C.P., Altenhofen, E., Ashworth, A., Brown, A., Kamau-Cheggeh, C., Curran, M., Evans, A., Floyd, R., Fowler, J., Garber, H., Hays, B., Kraemer, S., Lang, A., Mynhier, A., Samuels, A., Strohmaier, C., 2012. Ahrd Cyp1a2(−/−) mice show increased susceptibility to PCB-induced developmental neurotoxicity. Neurotoxicology 33, 1436–1442. https://doi.org/10.1016/j.neuro.2012.08.005

Curran, C.P., Vorhees, C.V., Williams, M.T., Genter, M.B., Miller, M.L., Nebert, D.W., 2011. In utero and lactational exposure to a complex mixture of polychlorinated biphenyls: toxicity in pups dependent on the Cyp1a2 and Ahr genotypes. Toxicol Sci 119, 189–208. https://doi.org/10.1093/toxsci/kfq314

Diliberto, J.J., Burgin, D., Birnbaum, L.S., 1997. Role of CYP1A2 in hepatic sequestration of dioxin: studies using CYP1A2 knock-out mice. Biochem Biophys Res Commun 236, 431–433. https://doi.org/10.1006/bbrc.1997.6973

Dix-Cooper, L., Eskenazi, B., Romero, C., Balmes, J., Smith, K.R., 2012. Neurodevelopmental performance among school age children in rural Guatemala is associated with prenatal and postnatal exposure to carbon monoxide, a marker for exposure to woodsmoke. Neurotoxicology 33, 246–254. https://doi.org/10.1016/j.neuro.2011.09.004

Dragin, N., Dalton, T.P., Miller, M.L., Shertzer, H.G., Nebert, D.W., 2006. For dioxin-induced birth defects, mouse or human CYP1A2 in maternal liver protects whereas mouse CYP1A1 and CYP1B1 are inconsequential. J Biol Chem 281, 18591–18600. https://doi.org/10.1074/jbc.M601159200

Duarte-Salles, T., Mendez, M.A., Meltzer, H.M., Alexander, J., Haugen, M., 2013. Dietary benzo(a)pyrene intake during pregnancy and birth weight: associations modified by vitamin C intakes in the Norwegian Mother and Child Cohort Study (MoBa). Environ Int 60, 217–223. https://doi.org/10.1016/j.envint.2013.08.016

Falcó, G., Domingo, J.L., Llobet, J.M., Teixidó, A., Casas, C., Müller, L., 2003. Polycyclic aromatic hydrocarbons in foods: human exposure through the diet in Catalonia, Spain. J Food Prot 66, 2325–2331. https://doi.org/10.4315/0362-028x-66.12.2325

Fuertes, E., Standl, M., Forns, J., Berdel, D., Garcia-Aymerich, J., Markevych, I., Schulte-Koerne, G., Sugiri, D., Schikowski, T., Tiesler, C.M.T., Heinrich, J., 2016. Traffic-related air pollution and hyperactivity/inattention, dyslexia and dyscalculia in adolescents of the German GINIplus and LISAplus birth cohorts. Environ Int 97, 85–92. https://doi.org/10.1016/j.envint.2016.10.017

Genkinger, J.M., Stigter, L., Jedrychowski, W., Huang, T.-J., Wang, S., Roen, E.L., Majewska, R., Kieltyka, A., Mroz, E., Perera, F.P., 2015. Prenatal polycyclic aromatic hydrocarbon (PAH) exposure, antioxidant levels and behavioral development of children ages 6-9. Environ Res 140, 136–144. https://doi.org/10.1016/j.envres.2015.03.017

IARC (International Agency for Research on Cancer. 2018. Monograph on Benzo[a]Pyrene.https://monographs.iarc.who.int/wp-content/uploads/2018/06/mono100F-14.pdf

ILAR (Institute for Laboratory Animal Research) 2011. Guide for the Care and Use of Laboratory Animals, 8th edition. National Research Council (US) Committee for the Update of the Guide for the Care and Use of Laboratory Animals. Washington (DC): National Academies Press (US); 2011.

Iyer, S., Perera, F., Zhang, B., Chanock, S., Wang, S., Tang, D., 2014. Significant interactions between maternal PAH exposure and haplotypes in candidate genes on B[a]P-DNA adducts in a NYC cohort of non-smoking African-American and Dominican mothers and newborns. Carcinogenesis 35, 69–75. https://doi.org/10.1093/carcin/bgt339

Iyer, S., Wang, Y., Xiong, W., Tang, D., Jedrychowski, W., Chanock, S., Wang, S., Stigter, L., Mróz, E., Perera, F., 2016. Significant interactions between maternal PAH exposure and single nucleotide polymorphisms in candidate genes on B[a]P-DNA adducts in a cohort of non-smoking Polish mothers and newborns. Carcinogenesis 37, 1110–1115. https://doi.org/10.1093/carcin/bgw090

Jedrychowski, W.A., Majewska, R., Spengler, J.D., Camann, D., Roen, E.L., Perera, F.P., 2017. Prenatal exposure to fine particles and polycyclic aromatic hydrocarbons and birth outcomes: a two-pollutant approach. Int Arch Occup Environ Health 90, 255–264. https://doi.org/10.1007/s00420-016-1192-9

Jedrychowski, W.A., Perera, F.P., Camann, D., Spengler, J., Butscher, M., Mroz, E., Majewska, R., Flak, E., Jacek, R., Sowa, A., 2015. Prenatal exposure to polycyclic aromatic hydrocarbons and cognitive dysfunction in children. Environ Sci Pollut Res Int 22, 3631–3639. https://doi.org/10.1007/s11356-014-3627-8

Korecka, A., Dona, A., Lahiri, S., Tett, A.J., Al-Asmakh, M., Braniste, V., D’Arienzo, R., Abbaspour, A., Reichardt, N., Fujii-Kuriyama, Y., Rafter, J., Narbad, A., Holmes, E., Nicholson, J., Arulampalam, V., Pettersson, S., 2016. Bidirectional communication between the Aryl hydrocarbon Receptor (AhR) and the microbiome tunes host metabolism. NPJ Biofilms Microbiomes 2, 16014. https://doi.org/10.1038/npjbiofilms.2016.14

Lam, J., Sutton, P., Kalkbrenner, A., Windham, G., Halladay, A., Koustas, E., Lawler, C., Davidson, L., Daniels, N., Newschaffer, C., Woodruff, T., 2016. A Systematic Review and Meta-Analysis of Multiple Airborne Pollutants and Autism Spectrum Disorder. PLoS One 11, e0161851. https://doi.org/10.1371/journal.pone.0161851

Lee, J., Kalia, V., Perera, F., Herbstman, J., Li, T., Nie, J., Qu, L.R., Yu, J., Tang, D., 2017. Prenatal airborne polycyclic aromatic hydrocarbon exposure, LINE1 methylation and child development in a Chinese cohort. Environ Int 99, 315–320. https://doi.org/10.1016/j.envint.2016.12.009

Li, C., Wang, J., Su, Q., Yang, K., Chen, C., Jiang, X., Han, T., Cheng, S., Mo, T., Zhang, R., Peng, B., Guo, Y., Baker, P.N., Tu, B., Xia, Y., 2018. Postnatal Subacute Benzo(a)Pyrene Exposure Caused Neurobehavioral Impairment and Metabolomic Changes of Cerebellum in the Early Adulthood Period of Sprague-Dawley Rats. Neurotox Res 33, 812–823. https://doi.org/10.1007/s12640-017-9832-8

Lin, Y.-C., Wu, C.-Y., Hu, C.-H., Pai, T.-W., Chen, Y.-R., Wang, W.-D., 2020. Integrated Hypoxia Signaling and Oxidative Stress in Developmental Neurotoxicity of Benzo[a]Pyrene in Zebrafish Embryos. Antioxidants (Basel) 9, E731. https://doi.org/10.3390/antiox9080731

Liu, S.-H., Wang, J.-H., Chuu, J.-J., Lin-Shiau, S.-Y., 2002. Alterations of motor nerve functions in animals exposed to motorcycle exhaust. J Toxicol Environ Health A 65, 803–812. https://doi.org/10.1080/00984100290071144

Lyu, Y., Ren, X.-K., Zhang, H.-F., Tian, F.-J., Mu, J.-B., Zheng, J.-P., 2020. Sub-chronic administration of benzo[a]pyrene disrupts hippocampal long-term potentiation via inhibiting CaMK II/PKC/PKA-ERK-CREB signaling in rats. Environ Toxicol 35, 961–970. https://doi.org/10.1002/tox.22932

Margolis, A.E., Herbstman, J.B., Davis, K.S., Thomas, V.K., Tang, D., Wang, Y., Wang, S., Perera, F.P., Peterson, B.S., Rauh, V.A., 2016. Longitudinal effects of prenatal exposure to air pollutants on self-regulatory capacities and social competence. J Child Psychol Psychiatry 57, 851–860. https://doi.org/10.1111/jcpp.12548

Margolis, A.E., Ramphal, B., Pagliaccio, D., Banker, S., Selmanovic, E., Thomas, L.V., Factor-Litvak, P., Perera, F., Peterson, B.S., Rundle, A., Herbstman, J.B., Goldsmith, J., Rauh, V., 2021. Prenatal exposure to air pollution is associated with childhood inhibitory control and adolescent academic achievement. Environ Res 202, 111570. https://doi.org/10.1016/j.envres.2021.111570

McCallister, M.M., Li, Z., Zhang, T., Ramesh, A., Clark, R.S., Maguire, M., Hutsell, B., Newland, M.C., Hood, D.B., 2016. Revealing Behavioral Learning Deficit Phenotypes Subsequent to In Utero Exposure to Benzo(a)pyrene. Toxicol Sci 149, 42–54. https://doi.org/10.1093/toxsci/kfv212

Min, J.-Y., Min, K.-B., 2017. Exposure to ambient PM10 and NO2 and the incidence of attention-deficit hyperactivity disorder in childhood. Environ Int 99, 221–227. https://doi.org/10.1016/j.envint.2016.11.022

Nebert, D.W., 2017. Aryl hydrocarbon receptor (AHR): “pioneer member” of the basic-helix/loop/helix per-Arnt-sim (bHLH/PAS) family of “sensors” of foreign and endogenous signals. Prog Lipid Res 67, 38–57. https://doi.org/10.1016/j.plipres.2017.06.001

Nebert, D.W., Shi, Z., Gálvez-Peralta, M., Uno, S., Dragin, N., 2013. Oral benzo[a]pyrene: understanding pharmacokinetics, detoxication, and consequences--Cyp1 knockout mouse lines as a paradigm. Mol Pharmacol 84, 304–313. https://doi.org/10.1124/mol.113.086637

Pagliaccio, D., Herbstman, J.B., Perera, F., Tang, D., Goldsmith, J., Peterson, B.S., Rauh, V., Margolis, A.E., 2020. Prenatal exposure to polycyclic aromatic hydrocarbons modifies the effects of early life stress on attention and Thought Problems in late childhood. J Child Psychol Psychiatry 61, 1253–1265. https://doi.org/10.1111/jcpp.13189

Patel, B., Das, S.K., Das, S., Das, L., Patri, M., 2016. Neonatal exposure to benzo[a]pyrene induces oxidative stress causing altered hippocampal cytomorphometry and behavior during early adolescence period of male Wistar rats. Int J Dev Neurosci 50, 7–15. https://doi.org/10.1016/j.ijdevneu.2016.01.006

Percie du Sert, N., Hurst, V., Ahluwalia, A., Alam, S., Avey, M.T., Baker, M., Browne, W.J., Clark, A., Cuthill, I.C., Dirnagl, U., Emerson, M., Garner, P., Holgate, S.T., Howells, D.W., Karp, N.A., Lazic, S.E., Lidster, K., MacCallum, C.J., Macleod, M., Pearl, E.J., Petersen, O.H., Rawle, F., Reynolds, P., Rooney, K., Sena, E.S., Silberberg, S.D., Steckler, T., Würbel, H., 2020. The ARRIVE guidelines 2.0: updated guidelines for reporting animal research. BMJ Open Sci 4, e100115. https://doi.org/10.1136/bmjos-2020-100115

Perera, F., Phillips, D.H., Wang, Y., Roen, E., Herbstman, J., Rauh, V., Wang, S., Tang, D., 2015. Prenatal exposure to polycyclic aromatic hydrocarbons/aromatics, BDNF and child development. Environ Res 142, 602–608. https://doi.org/10.1016/j.envres.2015.08.011

Perera, F., Weiland, K., Neidell, M., Wang, S., 2014. Prenatal exposure to airborne polycyclic aromatic hydrocarbons and IQ: estimated benefit of pollution reduction. J Public Health Policy 35, 327–336. https://doi.org/10.1057/jphp.2014.14

Perera, F.P., Chang, H., Tang, D., Roen, E.L., Herbstman, J., Margolis, A., Huang, T.-J., Miller, R.L., Wang, S., Rauh, V., 2014. Early-life exposure to polycyclic aromatic hydrocarbons and ADHD behavior problems. PLoS One 9, e111670. https://doi.org/10.1371/journal.pone.0111670

Perera, F.P., Tang, D., Wang, S., Vishnevetsky, J., Zhang, B., Diaz, D., Camann, D., Rauh, V., 2012. Prenatal polycyclic aromatic hydrocarbon (PAH) exposure and child behavior at age 6-7 years. Environ Health Perspect 120, 921–926. https://doi.org/10.1289/ehp.1104315

Perera, F.P., Wheelock, K., Wang, Y., Tang, D., Margolis, A.E., Badia, G., Cowell, W., Miller, R.L., Rauh, V., Wang, S., Herbstman, J.B., 2018. Combined effects of prenatal exposure to polycyclic aromatic hydrocarbons and material hardship on child ADHD behavior problems. Environ Res 160, 506–513. https://doi.org/10.1016/j.envres.2017.09.002

Peterson, B.S., Rauh, V.A., Bansal, R., Hao, X., Toth, Z., Nati, G., Walsh, K., Miller, R.L., Arias, F., Semanek, D., Perera, F., 2015. Effects of prenatal exposure to air pollutants (polycyclic aromatic hydrocarbons) on the development of brain white matter, cognition, and behavior in later childhood. JAMA Psychiatry 72, 531–540. https://doi.org/10.1001/jamapsychiatry.2015.57

Qiu, C., Peng, B., Cheng, S., Xia, Y., Tu, B., 2013. The effect of occupational exposure to benzo[a]pyrene on neurobehavioral function in coke oven workers. Am J Ind Med 56, 347–355. https://doi.org/10.1002/ajim.22119

Ribière, C., Peyret, P., Parisot, N., Darcha, C., Déchelotte, P.J., Barnich, N., Peyretaillade, E., Boucher, D., 2016. Oral exposure to environmental pollutant benzo[a]pyrene impacts the intestinal epithelium and induces gut microbial shifts in murine model. Sci Rep 6, 31027. https://doi.org/10.1038/srep31027

Rivas, I., Basagaña, X., Cirach, M., López-Vicente, M., Suades-González, E., Garcia-Esteban, R., Álvarez-Pedrerol, M., Dadvand, P., Sunyer, J., 2019. Association between Early Life Exposure to Air Pollution and Working Memory and Attention. Environ Health Perspect 127, 57002. https://doi.org/10.1289/EHP3169

Sentís, A., Sunyer, J., Dalmau-Bueno, A., Andiarena, A., Ballester, F., Cirach, M., Estarlich, M., Fernández-Somoano, A., Ibarluzea, J., Íñiguez, C., Lertxundi, A., Tardón, A., Nieuwenhuijsen, M., Vrijheid, M., Guxens, M., INMA Project, 2017. Prenatal and postnatal exposure to NO2 and child attentional function at 4-5years of age. Environ Int 106, 170–177. https://doi.org/10.1016/j.envint.2017.05.021

Singh, S.V., Benson, P.J., Hu, X., Pal, A., Xia, H., Srivastava, S.K., Awasthi, S., Zaren, H.A., Orchard, J.L., Awasthi, Y.C., 1998. Gender-related differences in susceptibility of A/J mouse to benzo[a]pyrene-induced pulmonary and forestomach tumorigenesis. Cancer Lett 128, 197–204. https://doi.org/10.1016/s0304-3835(98)00072-x

Uno, S., Dalton, T.P., Derkenne, S., Curran, C.P., Miller, M.L., Shertzer, H.G., Nebert, D.W., 2004. Oral exposure to benzo[a]pyrene in the mouse: detoxication by inducible cytochrome P450 is more important than metabolic activation. Mol Pharmacol 65, 1225–1237. https://doi.org/10.1124/mol.65.5.1225

Uno, S., Dalton, T.P., Dragin, N., Curran, C.P., Derkenne, S., Miller, M.L., Shertzer, H.G., Gonzalez, F.J., Nebert, D.W., 2006. Oral benzo[a]pyrene in Cyp1 knockout mouse lines: CYP1A1 important in detoxication, CYP1B1 metabolism required for immune damage independent of total-body burden and clearance rate. Mol Pharmacol 69, 1103–1114. https://doi.org/10.1124/mol.105.021501

Uno, S., Dragin, N., Miller, M.L., Dalton, T.P., Gonzalez, F.J., Nebert, D.W., 2008. Basal and inducible CYP1 mRNA quantitation and protein localization throughout the mouse gastrointestinal tract. Free Radic Biol Med 44, 570–583. https://doi.org/10.1016/j.freeradbiomed.2007.10.044

Vorhees, C.V., Williams, M.T., 2006. Morris water maze: procedures for assessing spatial and related forms of learning and memory. Nat Protoc 1, 848–858. https://doi.org/10.1038/nprot.2006.116

Wang, Y., Jiao, Y., Kong, Q., Zheng, F., Shao, L., Zhang, T., Jiang, D., Gao, X., 2021. Occurrence of polycyclic aromatic hydrocarbons in fried and grilled fish from Shandong China and health risk assessment. Environ Sci Pollut Res Int. https://doi.org/10.1007/s11356-021-13045-y

Zhang, H., Nie, J., Li, X., Niu, Q., 2013. Association of aryl hydrocarbon receptor gene polymorphism with the neurobehavioral function and autonomic nervous system function changes induced by benzo[a]pyrene exposure in coke oven workers. J Occup Environ Med 55, 265–271. https://doi.org/10.1097/JOM.0b013e318278272f

Zhou, S.-F., Wang, B., Yang, L.-P., Liu, J.-P., 2010. Structure, function, regulation and polymorphism and the clinical significance of human cytochrome P450 1A2. Drug Metab Rev 42, 268–354. https://doi.org/10.3109/03602530903286476

